# Efficacy of SGLT2 Inhibitors in Pulmonary Arterial Hypertension: A Systematic Review and Meta-Analysis of Preclinical Studies

**DOI:** 10.64898/2026.03.18.712480

**Authors:** Fares Qubbaj, Ahmad Saeed, Osama Younis, Nada Al-Awamleh, Zeid Al-Sharif, Qais Shaban, Samia Sulaiman, Ahmad Turk

**Author notes:** **Correspondence**: Fares Qubbaj, School of Medicine, University of Jordan, Amman, Jordan.

## Abstract

**Background:** Pulmonary arterial hypertension (PAH) is a progressive disease marked by vascular remodeling, elevated pulmonary pressures, and right ventricular failure. Current therapies are mainly vasodilatory, underscoring the need for treatments targeting additional pathways. Sodium–glucose cotransporter-2 (SGLT2) inhibitors, initially used for diabetes, have demonstrated cardiovascular benefits.

**Aims:** This systematic review and meta-analysis evaluated the effects of SGLT2 inhibitors in animal models of PAH, focusing on pulmonary hemodynamics and right ventricular function.

**Methods:** PubMed, Embase, Web of Science, and Scopus were searched for preclinical studies reporting mean pulmonary artery pressure (mPAP), right ventricular systolic pressure (RVSP), right ventricular hypertrophy index (RV/LV+S), tricuspid annular plane systolic excursion (TAPSE), or pulmonary artery acceleration time (PAAT). Random-effects meta-analyses were performed using R.

**Results:** Nine studies were included. SGLT2 inhibitors were significantly associated with lower mPAP (WMD −9.79 mmHg), RVSP (WMD −14.81 mmHg), and RV/LV+S (WMD −0.10). They were also associated with higher indices of right ventricular function, including TAPSE (WMD 0.53 mm) and PAAT (WMD 6.39 ms).

**Conclusion:** In preclinical models of PAH, SGLT2 inhibitor treatment was associated with favorable hemodynamic and structural parameters. Further research is needed to clarify their translational potential and long-term safety.

## 1.0 Introduction

Pulmonary arterial hypertension (PAH), also classified as group 1 pulmonary hypertension, is a chronic, progressive disease characterized by elevated pressure in the pulmonary arteries due to vascular remodeling and increased resistance to blood flow [1,2]. PAH predominantly affects younger individuals, imposing a severe health and socioeconomic burden, with a global incidence of approximately 5 adults per million per year and an associated prevalence of 15 per million adults [3].

The pathophysiology of PAH is multifaceted and is linked to its underlying etiology. Despite the variability in causes, common pathological features include increased pulmonary vascular resistance, endothelial dysfunction, vascular remodeling due to the chronic inflammation, and metabolic dysfunction, all of which contribute to right ventricular failure and death. These processes are driven by disruptions in nitric oxide (NO), prostacyclin (PGI2), thromboxane A2 (TXA2), and endothelin-1 (ET-1) signaling pathways, which form the foundation of PAH pathogenesis and targeted therapy [4].

Current PAH therapies target key pathophysiological mechanisms using phosphodiesterase-5 inhibitors, guanylate cyclase stimulators, prostacyclin analogues, receptor agonists, and endothelin receptor antagonists—alone or in combination. While these treatments have improved exercise capacity and quality of life, and in some cases delayed disease progression, their impact on long-term mortality remains limited, underscoring the urgent need for novel therapeutic strategies [5].

However, with the advent of Sodium-Glucose Cotransporter 2 inhibitors (SGLT2i) emerges the potential for these medications to be explored as adjuncts to existing treatment regimens for PAH. Their proven benefits in heart failure, particularly in attenuating cardiac remodeling, suggest potential cardioprotective effects in PAH [6]. Furthermore, by modulating metabolic dysfunction—a key driver of pulmonary vascular remodeling—they represent a promising therapeutic avenue [7].

The literature on SGLT2i in pulmonary arterial hypertension (PAH) shows notable inconsistencies. Li et al. found that dapagliflozin had no protective effect on right ventricular remodeling or function in PAH rats [8], whereas Wu et al. reported significant improvements in both parameters [9].

Amid these contradictions, this systematic review and meta-analysis aims to synthesize and quantitatively assess preclinical evidence on the effects of SGLT2i in experimental animal models of pulmonary hypertension. As current evidence stems mainly from preclinical animal studies, our analysis addresses a critical preclinical gap and provides hypothesis-generating insights to inform future mechanistic and translational studies, given the absence of completed human clinical trials [10].

## 2.0 Methodology

This systematic review and meta-analysis followed the Systematic Review Centre for Laboratory Animal Experimentation (SYRCLE) guidelines [11]. The study protocol was registered in the PROSPERO database [12] (CRD42024627817), which contains a detailed account of our methods. A complete PRISMA checklist is available in the supplementary material (S1).

### 2.1 Search Strategy

A literature search was performed on Nov 10th, 2024, across four electronic databases: PubMed, Web of Science, Scopus, and Embase. The search included terms like “Animal Model,” “Preclinical,” “Mice,” “SGLT2 Inhibitors,” “empagliflozin,” “Pulmonary Arterial Hypertension,” and “Right Ventricular Dysfunction.” Full search strings are in the supplementary material (S2).

### 2.2 Study Selection

Two independent reviewers (QS and NA) screened the articles using blinded review via Rayyan.ai [13]. Screening began with titles and abstracts, followed by full-text review based on predefined inclusion and exclusion criteria. Discrepancies were resolved through discussion or consultation with the primary investigator (FQ).

The predetermined inclusion criteria used in both phases were (a) Studies published in English, (b) Experimental animal studies such as rats, mice, pigs, etc., (c) Group 1 pulmonary hypertension, (d) SGLT2i therapy, (e) Studies with control group (f) Quantitative reporting of cardiovascular outcomes or hemodynamic measurements. Exclusion criteria were (a) Studies published in a language other than English, (b) Non-primary research articles such as reviews, perspective papers, or letters to the editor, (c) Studies with incorrect designs or interventions that did not align with the objectives of this review, (d) in vitro/in silico studies, (e) Lack of SGLT2i intervention, and (f) studies presenting only qualitative data on cardiovascular outcomes or hemodynamic measurements.

### 2.3 Quality Assessment

Two reviewers (NA and ZA) independently assessed the risk of bias using SYRCLE’s Risk of Bias tool [11], which evaluates 10 domains across six bias categories. Disagreements were resolved by consensus or by consulting the primary investigator (FQ).

### 2.4 Data Extraction

Extracted data included study characteristics (authors, year of publication), experimental details (mode of PAH induction, animal model, sample size and grouping, duration of treatment, regimen of treatment including route of delivery, dose, and frequency), and outcome measures (mean ± SD for drug and placebo groups).

For studies reporting the standard error of the mean (SEM) instead of standard deviation (SD), SD was calculated using the formula *SD* = *SEM* × √*n*, where n is the sample size. In studies reporting two doses of the drug, a single dose was selected according to predefined criteria to maintain comparability with doses reported in other studies; specifically, the lower dose was used when it more closely aligned with the dosing range of the included literature.

Data were extracted from text, tables, supplementary files, or graphs. For graphical data, the trained authors utilized PlotDigitizer to accurately extract values [14]. For error bars in figures representing measures of variation (SD or SEM) shown in one direction from the mean, symmetry above and below the mean was assumed.

Missing data were addressed by contacting study authors directly for clarification or additional information. Data extraction was conducted independently by two reviewers (AS and QS), with discrepancies resolved by consensus with the primary investigator (FQ).

### 2.5 Meta-Analysis

A meta-analysis was conducted to evaluate the effect of SGLT2i on PAH-related outcomes. The primary measure of effect was the mean difference between drug-treated and placebo groups, reported with 95% confidence intervals (95%CI). To address potential dependency bias, outcomes reported more than once within the same study were aggregated into a single effect size per study. Aggregation was performed using the aggregate.escalc() function in the metafor package, which applies inverse-variance weighting to compute a combined effect size. This approach ensured that each study contributed only one independent effect size to the meta-analysis.

The overall effect was calculated as a weighted mean difference (WMD) across studies. A random-effects model, using the DerSimonian and Laird method, was used for pooling effect sizes, accounting for variability between studies. Study weighting was based on the inverse variance method. Heterogeneity was assessed using Higgins and Thompson’s I² statistic to quantify the degree of inconsistency across studies and τ² to estimate between-study variance.

Subgroup analyses were performed to examine treatment effects based on the specific SGLT2i drug (dapagliflozin, canagliflozin, empagliflozin) and animal model. For the animal model subgroup analysis, the lack of statistical significance between groups supported the justification for aggregating outcomes in studies reporting the same outcome twice, once per model. Leave one out sensitivity analysis was conducted to test the robustness of the results and identify potential sources of heterogeneity. Publication bias was not assessed, as none of the outcomes included data from 10 or more studies, consistent with standard recommendations [15].

All statistical analyses were performed using R version 4.4.2 (R Foundation for Statistical Computing, Vienna, Austria) within the RStudio environment (RStudio version 2024.04.1+748, Posit Software, 2024). Meta-analyses were conducted using the “meta” and “metafor” packages [16,17], and results were presented as forest plots. Statistical significance was set at P<0.05.

## 3.0 Results

### 3.1 Study selection

A total of 1,827 studies were retrieved from PubMed, Web of Science, Embase, and Scopus. After removing 828 duplicates, 999 studies were screened. Of these, 984 were excluded based on predefined criteria, leaving 15 for full-text review. Six studies were excluded (2 non-PAH, 4 inappropriate induction), resulting in 9 included studies. A PRISMA flow diagram (Figure 1) outlines the selection process.

**Figure 1:**
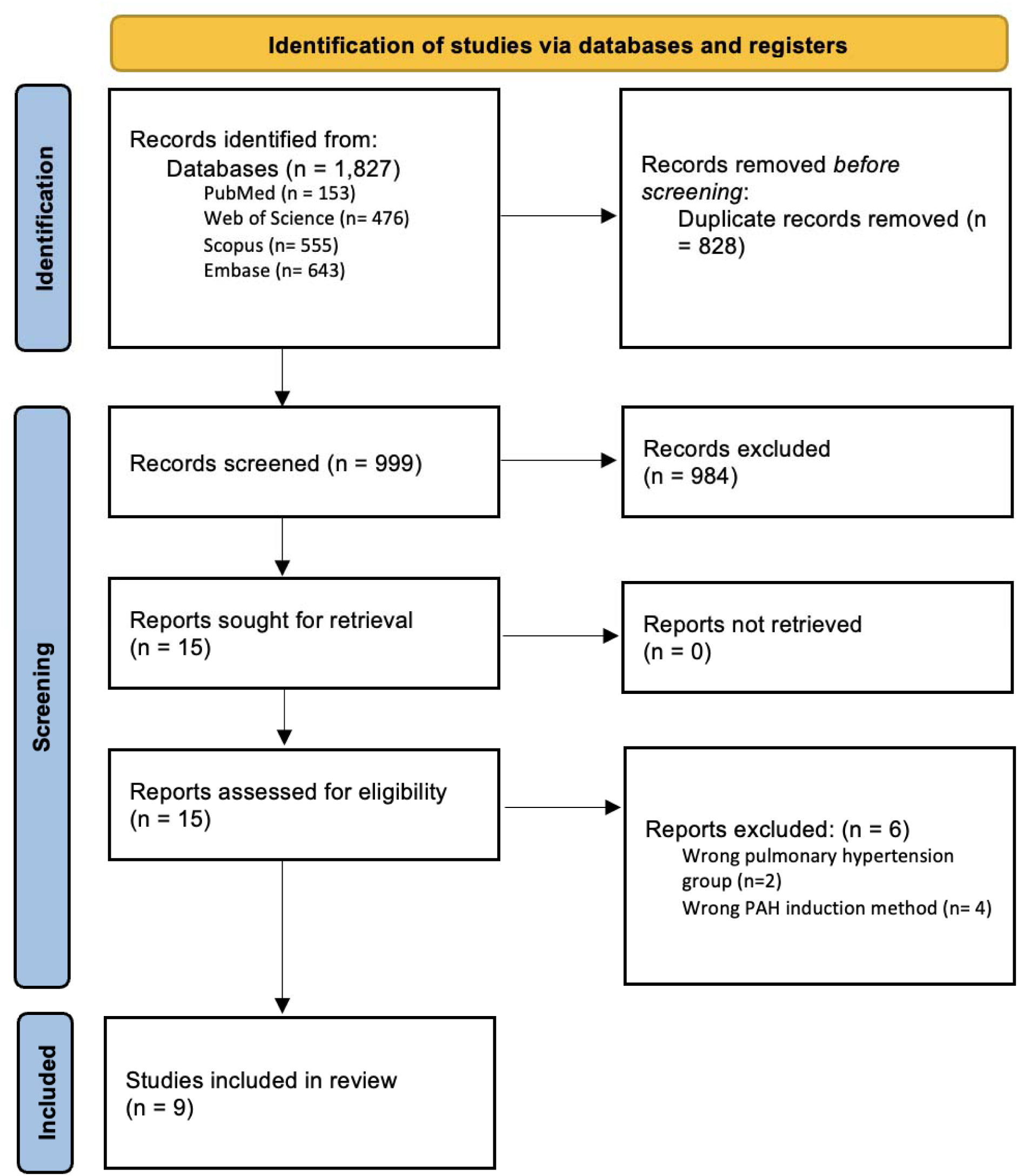
PRISMA flowchart

### 3.2 Characteristics of included studies

Nine studies, published between 2020 and 2024, using rodent models were included. Eight used rats; one (Li et al., 2024[18]) included both rats and mice. PAH was induced using monocrotaline, Sugen/hypoxia, or Sugen/hypoxia/normoxia [19]. Dapagliflozin was most commonly used. Outcomes assessed included mPAP, RVSP, RV/LV+S, TAPSE, and PAAT. Study characteristics are summarized in Table 1.

**Table:** Summary of SGLT2 inhibitor (SGLT2i) studies in pulmonary arterial hypertension included in this systematic review and meta-analysis. Abbreviations: MCT, monocrotaline; SC, subcutaneous; IP, intraperitoneal; Su/Hx, Sugen5416 + hypoxia; Su/Hx/Nx, Sugen5416 + hypoxia+ Normoxia. The table summarizes key details such as study ID, drug regimen, sample size, experimental groupings, animal models, and methods of pulmonary hypertension induction across all included studies.

| StudyID | Drug | Regimen<br>(Dose, Route,<br>Duration) | Treatment timing<br>relative to PH<br>induction | Sample<br>Size<br>(n/group) | Groupings | Animal<br>Model | Mode of PH<br>Induction |
| --- | --- | --- | --- | --- | --- | --- | --- |
| Dai et al.,<br>2022 | Dapagliflozin | 60 mg/L, free<br>drinking, 4<br>weeks | Day 1 | 16/group | MCT+vehicle,<br>MCT+Dapagliflozin | Sprague-Dawley rats<br>(300–350 g) | MCT (60 mg/kg,<br>IP) |
| Wu et al.,<br>2022 | Dapagliflozin | 1 mg/kg/day,<br>oral gavage, 35<br>days | Day 1 | 8/group | MCT+vehicle,<br>MCT+Dapagliflozin | Sprague-Dawley rats<br>(220–280 g) | MCT (60 mg/kg,<br>IP) |
| Tang et al.,<br>2022 | Dapagliflozin | 1 mg/kg/day,<br>oral gavage, 3<br>weeks | Day 1 | 10/group | MCT+vehicle,<br>MCT+Dapagliflozin | Sprague-Dawley rats<br>(160–180 g) | MCT (60 mg/kg,<br>IP) |
| Qin et al.,<br>2022 | Dapagliflozin | 60 mg/L, free<br>drinking, 4<br>weeks | Day 0 (Same Day) | 20/group | MCT+vehicle,<br>MCT+Dapagliflozin | Sprague-Dawley rats<br>(300–350 g) | MCT (60 mg/kg,<br>IP) |
| Liu et al.,<br>2024 | Dapagliflozin | 0.5<br>mg/kg/day,<br>oral gavage, 5<br>weeks | Day 28(MCT); Day<br>35(Sugen/Hypoxia) | 6/group | MCT+ vehicle,<br>MCT+Dapagliflozin; Su/Hx+<br>vehicle, Su/Hx+Dapagliflozin | Sprague-<br>Dawley rats<br>(250–300 g) | MCT (60 mg/kg,<br>IP); Sugen5416<br>(20 mg/kg, SC) +<br>hypoxia (FiO<sub>2</sub><br>10%) |
| Li et al., 2021 | Dapagliflozin | 1.5<br>mg/kg/day,<br>oral gavage, 5<br>weeks | Day 1 | 15/group | MCT+Vehicle,<br>MCT+Dapagliflozin | Sprague-Dawley rats<br>(180–200 g) | MCT (60 mg/kg,<br>SC) |
| Chen et al.,<br>2024 | Canagliflozin | 30 mg/kg/day,<br>oral gavage, 4<br>weeks | Day 1 | 8/group | MCT+vehicle,<br>MCT+Canagliflozin | Sprague-Dawley rats<br>(200–230 g) | MCT (40 mg/kg,<br>SC) |
| Li et al., 2024 | Canagliflozin | 10 mg/kg/day,<br>oral gavage, 21<br>days | Day 1 | 6 mice<br>/group | Su/Hx+vehicle, Su/Hx +<br>Canagliflozin;<br>Su/Hx/Nx+vehicle, | C57BL/6J<br>mice (20–25<br>g) | Sugen5416 (20<br>mg/kg, SC) +<br>hypoxia chamber |
|  |  |  |  |  | Su/Hx/Nx + Canagliflozin |  | (FiO<sub>2</sub> 10%) |
|  |  | 5 mg/kg/day,<br>oral gavage, 21<br>days | Day 1 | 8 rats<br>/group | MCT+vehicle, MCT +<br>Canagliflozin;<br>Su/Hx+vehicle, Su/Hx +<br>Canagliflozin | Sprague-Dawley rats<br>(200–250 g) | MCT (60 mg/kg,<br>IP); Sugen5416<br>(20 mg/kg, SC) +<br>hypoxia |
| **Chowdhury**<br>**et al., 2020** | Empagliflozin | 10 mg/kg/day,<br>oral gavage, 28<br>days | Day 1 | 16/group | MCT+Vehicle,<br>MCT+Empagliflozin | Sprague-Dawley rats<br>(220–250 g) | MCT (60 mg/kg,<br>IP) |

### 3.3 Risk of bias

Risk of bias was assessed using SYRCLE’s RoB tool. Random sequence generation, random housing, blinding of outcome assessment, and selective outcome reporting showed low risk. Baseline characteristics were consistently reported. Allocation concealment, random outcome assessment, incomplete outcome data, and other domains had mostly unclear risk. High risk was noted in “Blinding of Participants and Personnel” for Li et al. and Qin et al., and in “Blinding of Outcome Assessment” for Li et al. Results are visualized in Figure 2; detailed judgments are in Supplementary Material (S3).

**Figure 2:**
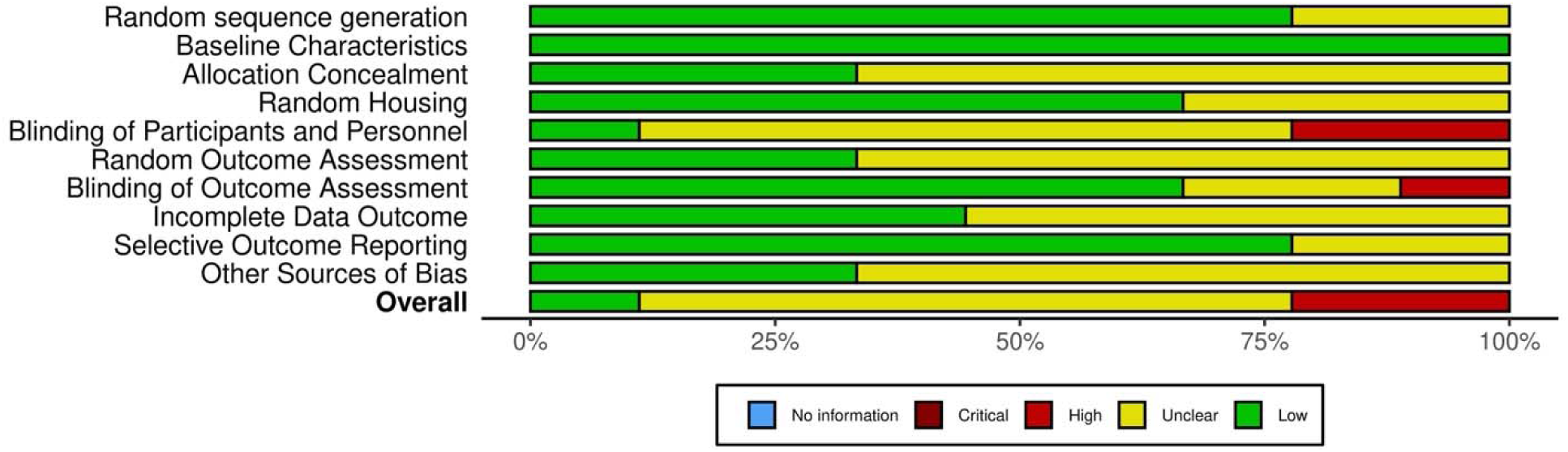
Quality assessment of the studies included in the meta-analysis using the SYRCLE risk of bias tool. The left panel presents the 10 domains that the SYRCLE assesses studies against, each falling under one of the 6 tested biases. Red: high risk of bias, Green: low risk of bias. Yellow: unclear risk of bias.

### 3.4 Mean pulmonary artery pressure (mPAP)

Meta-analysis of 3 studies showed SGLT2i significantly reduced mPAP compared to controls [9,20,21], with a WMD of −9.79 mmHg (95% CI: −14.67, −4.92; p < 0.0001) and no heterogeneity (I² = 0%, τ² = 0, p = 0.5146) (Figure 3a). Subgroup analysis by drug showed dapagliflozin had a WMD of −11.61 mmHg (95% CI: −17.39, −5.84), while empagliflozin had a WMD of −5.28 mmHg (95% CI: −14.37, 3.8), with no significant difference between drugs (p = 0.2493) (Figure 3b). Sensitivity analysis confirmed no single study influenced the overall effect (p < 0.0001) (Figure 3c).

**Figure 3.**
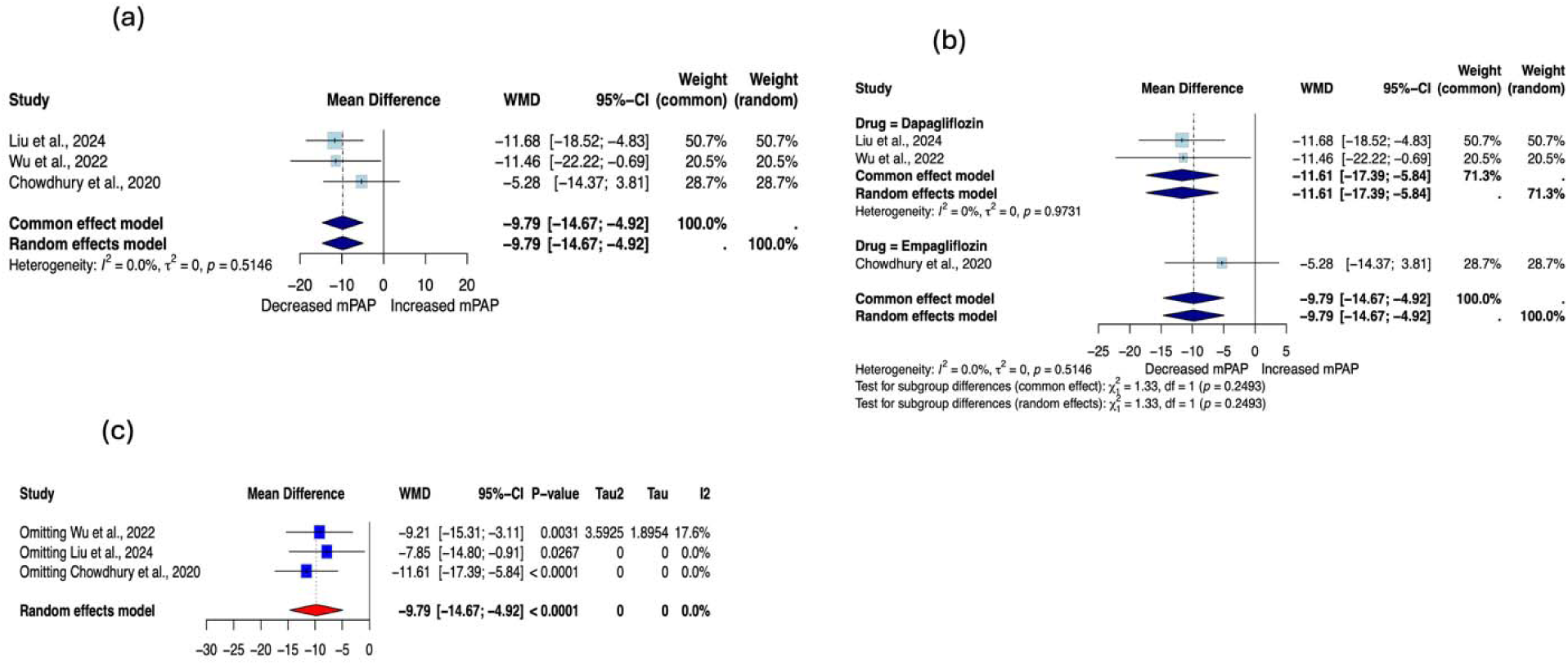
Meta-analysis of mPAP in preclinical PAH models. (A) Overall effect estimate for mPAP across all included studies. (B) Subgroup analysis based on the specific SGLT2 inhibitor administered. (C) Leave-one-out analysis (LOOA) assessing the influence of individual studies on the pooled effect estimate.

### 3.5 Right ventricular systolic pressure (RVSP)

Six studies showed significant RVSP reduction with SGLT2i[8,18,20–23], with a WMD of −14.81 mmHg (95% CI: −19.50, −10.12; p < 0.0001) and moderate heterogeneity (I² = 63.4%, τ² = 21.79, p = 0.0117) (Figure 4a). Animal model subgroup analysis showed no significant differences (p = 0.3071), supporting pooling of studies using different animal models for the meta-analysis (Figure 4b). Drug-specific analysis showed empagliflozin had a WMD of −21.78 mmHg (95% CI: −26.40, −17.16) and dapagliflozin had a WMD of −13.33 mmHg (95% CI: −24.38, −2.29), with a significant subgroup difference (p = 0.0338)(Figure 4c). Sensitivity analysis confirmed robustness (p < 0.0001) (Figure 4d).

**Figure 4.**
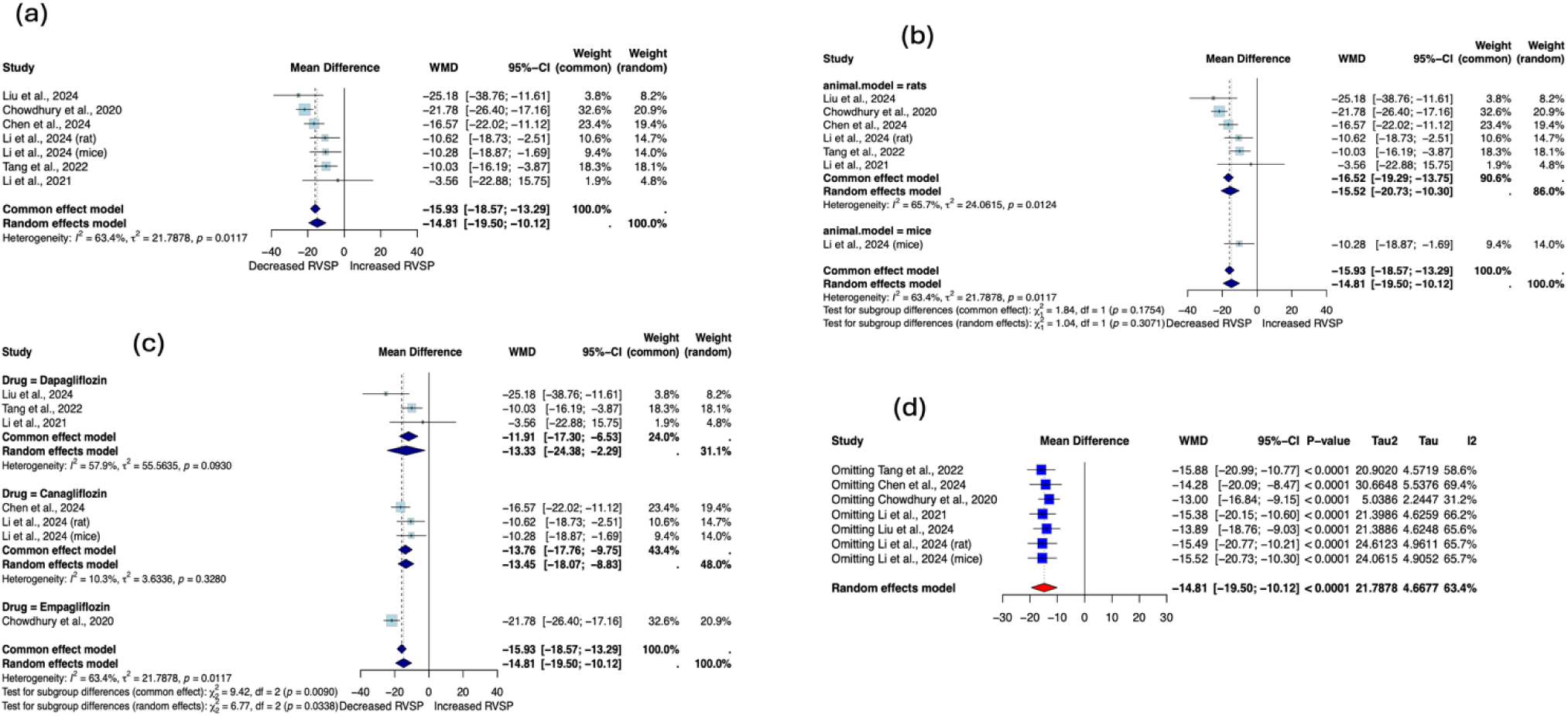
Meta-analysis of RVSP in preclinical PAH models. (A) Overall effect estimate for RVSP across all included studies. (B) Subgroup analysis based on animal model used. (C) Subgroup analysis based on the specific SGLT2 inhibitor administered. (D) Leave-one-out analysis (LOOA) assessing the influence of individual studies on the pooled effect estimate.

### 3.6 Right ventricular hypertrophy index (RV/LV+S)

Eight studies showed a significant reduction in RV/LV+S with SGLT2i therapy, with a [8,9,18,20–22,24,25]WMD of −0.10 (95% CI: −0.12, −0.07; p < 0.0001) and no heterogeneity (I² = 0%, τ² = 0, p = 0.4779) (Figure 5a). No significant difference was found by animal model (p = 0.6360), supporting pooled analysis (Figure 5b). Drug subgroup WMDs were −0.10 (95% CI: −0.14, −0.06) for dapagliflozin, −0.10 (95% CI: −0.30, −0.09) for empagliflozin, and −0.09 (95% CI: −0.13, −0.05) for canagliflozin, with no significant inter-drug difference (p = 0.9467) (Figure 5c). Sensitivity analysis confirmed results (p < 0.0001) (Figure 5d).

**Figure 5.**
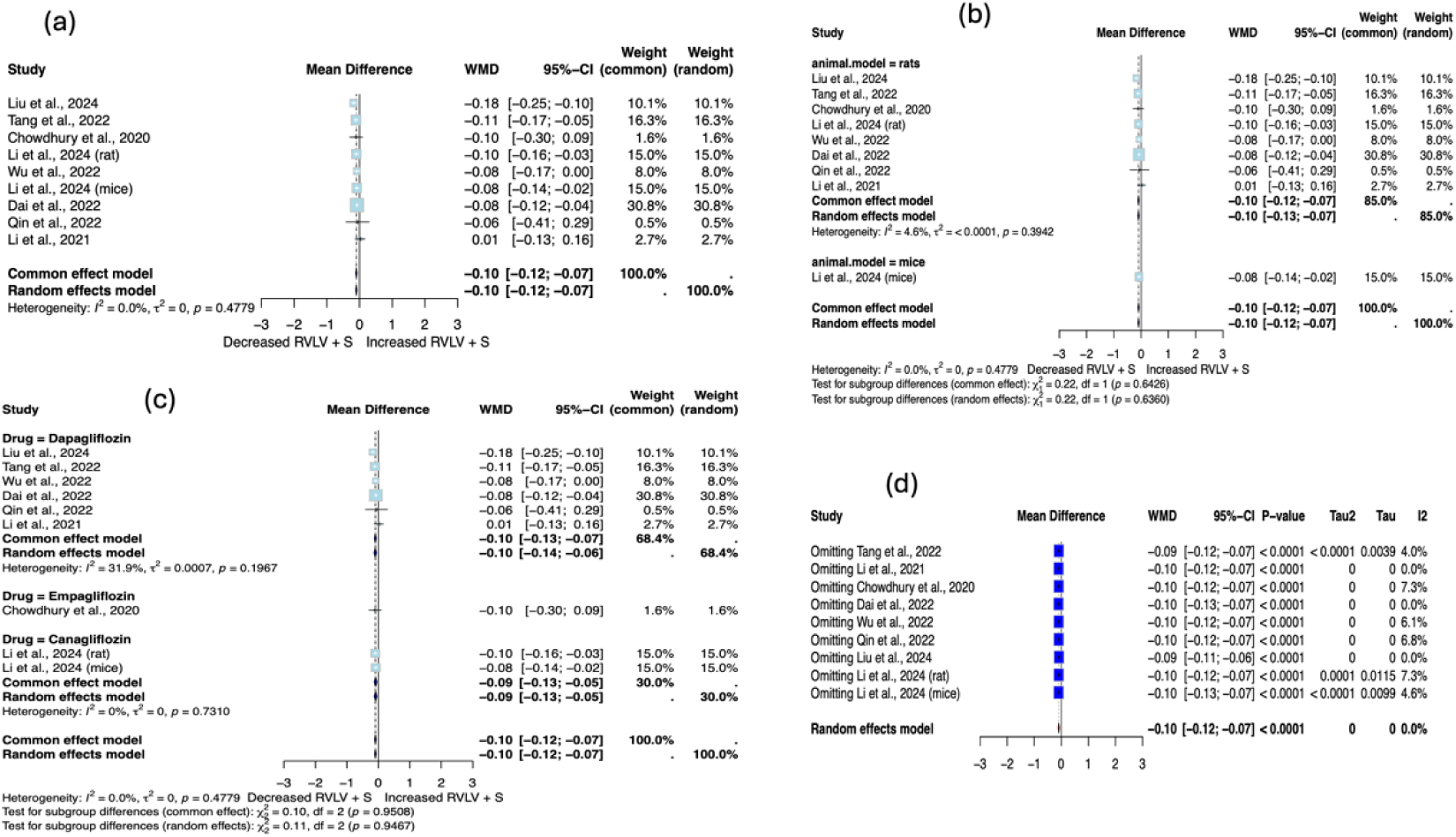
Meta-analysis of RV/LV+S in preclinical PAH models. (A) Overall effect estimate for RV/LV+S across all included studies. (B) Subgroup analysis based on animal model used. (C) Subgroup analysis based on the specific SGLT2 inhibitor administered. (D) Leave-one-out analysis (LOOA) assessing the influence of individual studies on the pooled effect estimate.

### 3.7 Tricuspid annular plane systolic excursion (TAPSE)

Six studies showed TAPSE was significantly increased by SGLT2i[8,9,20,23–25], with a WMD of 0.53 mm (95% CI: 0.22, 0.85; p = 0.0008) and moderate heterogeneity (I² = 63.4%, τ² = 0.0843, p = 0.0180) (Figure 6a). Drug subgroup WMDs were 0.78 mm (95 CI: 0.24, 1.32) for canagliflozin and 0.49 mm (95% CI: 0.12, 0.85) for dapagliflozin, with no significant inter-drug difference (p = 0.3754) (Figure 6b). Sensitivity analysis showed robust results (p = 0.0008) (Figure 6c).

**Figure 6.**
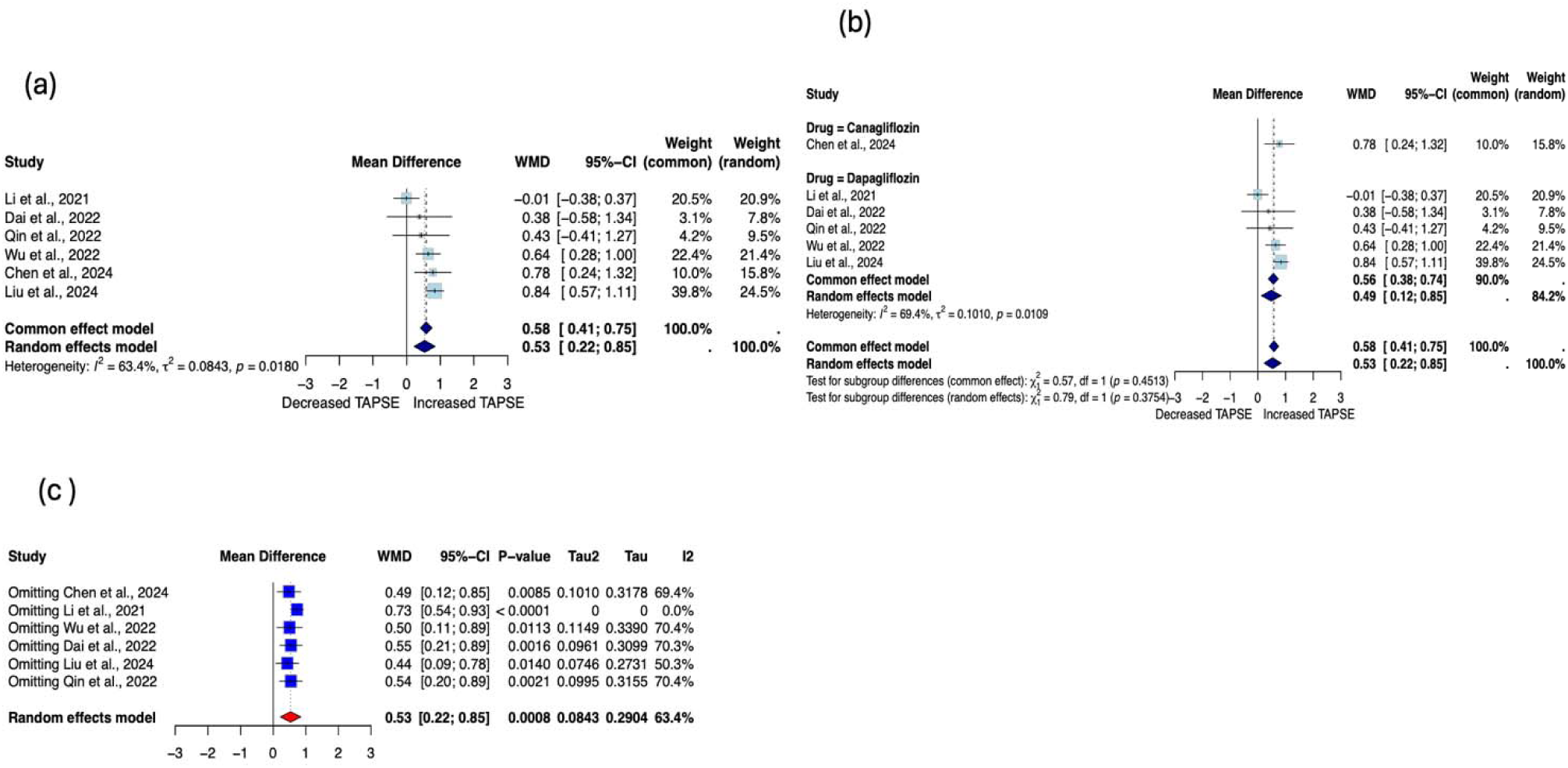
Meta-analysis of TAPSE in preclinical PAH models. (A) Overall effect estimate for TAPSE across all included studies. (B) Subgroup analysis based on the specific SGLT2 inhibitor administered. (C) Leave-one-out analysis (LOOA) assessing the influence of individual studies on the pooled effect estimate.

### 3.8 Pulmonary artery acceleration time (PAAT)

Seven studies showed a significant PAAT increase with SGLT2i therapy [8,9,18,21,23–25], with a WMD of 6.39 ms (95% CI: 5.46, 7.32; p < 0.0001) and low heterogeneity (I² = 8.9%, τ² = 0, p = 0.3611) (Figure 7a). No significant difference by animal model was found (p = 0.5853), supporting pooled analysis (Figure 7b). Drug subgroup WMDs were 2.75 ms (95% CI: −2.22, 7.71) for dapagliflozin, 6.39 ms (95% CI: 5.37, 7.41) for empagliflozin, and 7.96 ms (95% CI: 4.07, 11.85) for canagliflozin, with no significant differences between drugs (p = 0.2611) (Figure 7c). Sensitivity analysis confirmed a consistent effect (p < 0.0001) (Figure 7d).

**Figure 7.**
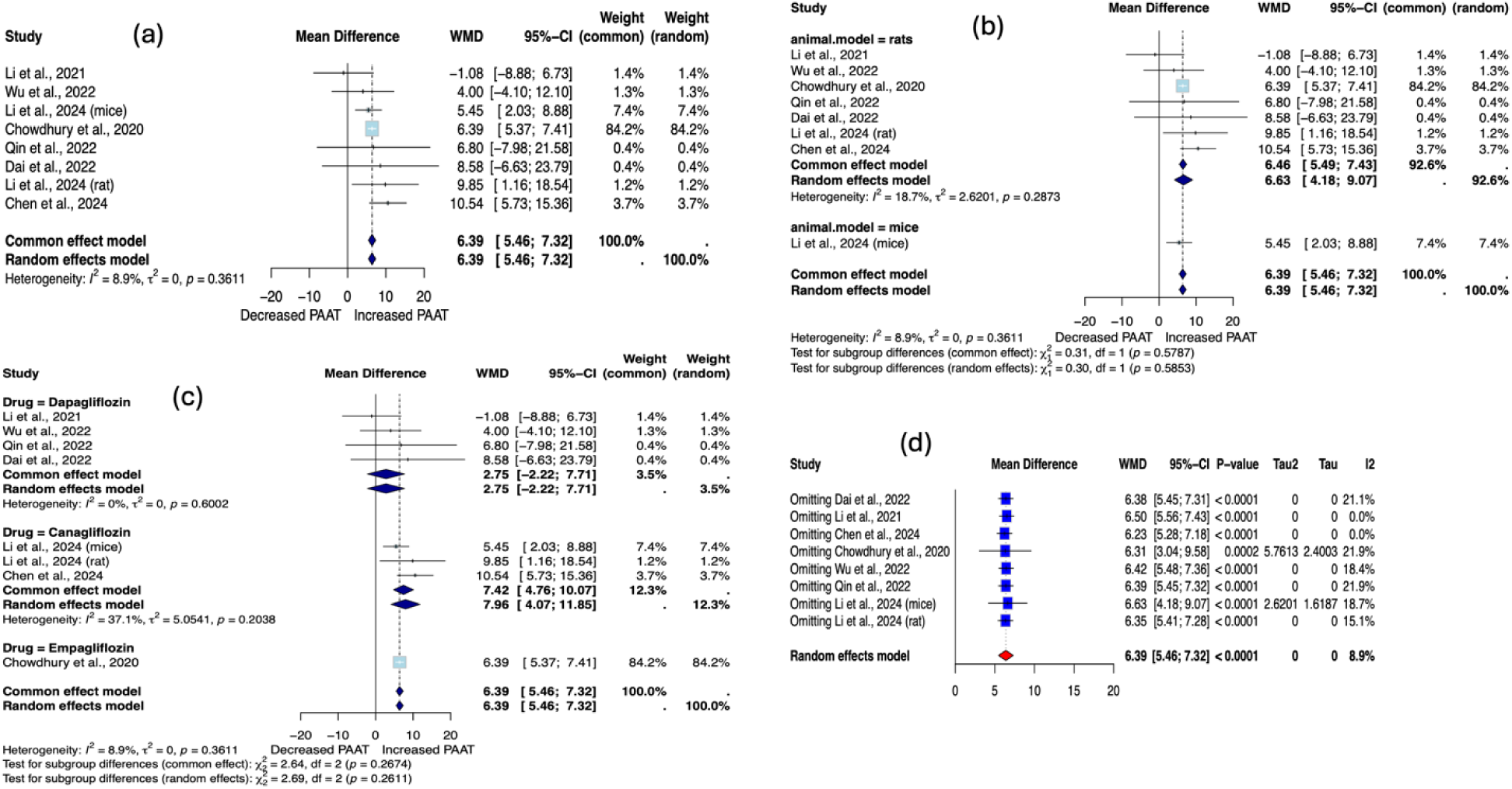
Meta-analysis of PAAT in preclinical PAH models. (A) Overall effect estimate for PAAT across all included studies. (B) Subgroup analysis based on animal model used. (C) Subgroup analysis based on the specific SGLT2 inhibitor administered. (D) Leave-one-out analysis (LOOA) assessing the influence of individual studies on the pooled effect estimate.

## 4.0 Discussion

### 4.1 Summary of findings

Our findings demonstrate that SGLT2i are associated with significant reductions in mPAP, RV/LV+S, and RVSP, as well as improvement in markers of right ventricular function, PAAT and TAPSE, in preclinical models of PAH. The efficacy of SGLT2i was consistent upon subgroup analysis across animal models (rats vs. mice) as well as drug types (empagliflozin, dapagliflozin, canagliflozin).

### 4.2 Mechanism of action of SGLT2 inhibitors in PAH

The beneficial effects of SGLT2i in experimental animal models of PAH have been proposed to stem from multifaceted mechanisms, as highlighted by Luo et al. [26]. These include mitigating endothelial dysfunction, central to PAH pathophysiology, mainly by reducing glycemic variability and oxidative stress. Endothelial dysfunction is a major contributor to elevated mPAP and RVSP, and its amelioration can directly reduce vascular resistance and improve pulmonary arterial compliance, since T1R/NADPH oxidase/SGLT1 and 2 pathways promote endothelial dysfunction. The study has also highlighted that empagliflozin and dapagliflozin both restore nitric oxide’s bioavailability through the inhibition of reactive oxygen species, thus improving endothelial function [26]. They also inhibit endothelial-to-mesenchymal transition (EndoMT), a key process in PAH, and attenuate vascular neointimal hyperplasia via the TAK-1/NF-kB pathway, as demonstrated in murine carotid models [27].

Furthermore, they reduce pulmonary artery smooth muscle cell (PASMC) proliferation, type 1 collagen deposition, and promote apoptosis, thereby alleviating vascular remodeling [26]. Through minimizing the proportion of proliferating cell nuclear antigen positive cells in the pulmonary artery and attenuating the expression of Cx43 and CD31 in the RV, it is believed that dapagliflozin inhibits PASMC proliferation in such mechanism, while other agents such as tofogliflozin decrease PASMC migration instead. These findings align with our findings demonstrating reductions in mPAP and improved pulmonary arterial load, as reflected by increased PAAT [26,27].

Anti-inflammatory actions, including IL-6 and TNF-α downregulation, further reduce vascular remodeling and RV pressure overload, consistent with RVSP improvements and enhanced right ventricular function [28]. Additionally, ion channel modulation stabilizes vascular tone and contractility, supporting better systolic function, as reflected by increased TAPSE.

Collectively, these experimentally observed mechanisms suggest that SGLT2i may influence multiple components of pulmonary vascular remodeling and right ventricular adaptation in animal models, potentially contributing to the hemodynamic and functional improvements reported in individual studies. Additionally, histological findings of dapagliflozin and empagliflozin are observed in numerous studies. For example, Chowdhury et al. have found increased apoptosis and decreased proliferation in empagliflozin-treated lung blood vessels in monocrotaline PAH rats, indicating that empagliflozin prevents adverse pulmonary arteriole remodeling. Their study also revealed significantly reduced medial wall thickening and decreased muscularization of pulmonary arterioles in the group with empagliflozin treatment [21].

### 4.3 Comparison to previous studies

Over the recent years, SGLT2i have gained widespread recognition for their indirect cardioprotective effects, independent of their anti-diabetic role. Landmark trials like “EMPEROR Reduced” and “DAPA-HF” have shown that SGLT2i reduced mortality and cardiac hospitalizations in left ventricular failure patients [29,30]. However, the left and right ventricles differ significantly in their anatomy, function, and mechanisms of failure, and hence the therapeutic effects of SGLT2i on pulmonary vascular remodeling and subsequently the right ventricle, cannot be extrapolated from existing research focused on left heart failure [31].

This distinction underscores the importance of investigating the cardioprotective mechanisms of SGLT2i in the context of pulmonary arterial hypertension and right heart function, where pulmonary vascular remodeling plays a central role in disease progression and outcomes. In this regard, our findings align with prior evidence, such as the work by Tan et al., which demonstrated the ability of SGLT2i to mitigate PAH progression [27]. However, only one study included in our analysis has shown that dapagliflozin has no cardioprotective effect, despite showing similar baseline characteristics and intervention regimen to the other studies [8]. This suggests that subtle differences in laboratory conditions, such as variable animal handling or environmental stressors, could have played an unrecognized role. To better understand these differences, studies should minimize and report the variability in baseline characteristics and experimental conditions.

SGLT2i’s effects in PAH can also be contextualized against other drug classes commonly used in PAH management, such as phosphodiesterase-5 (PDE5) inhibitors and endothelin receptor antagonists (ERAs). While PDE5 inhibitors and ERAs mainly work by promoting vasodilation and preventing endothelin mediated vasoconstriction respectively, SGLT2i have shown their ability to address inflammation, vascular remodeling, and oxidative stress as well [32,33]. Theoretically, these proposed mechanisms make SGLT2i great candidates for potential complementary use with established PAH treatments.

To our best knowledge, only one study to date has explored the combination of SGLT2i with a specific PAH therapy, namely dapagliflozin with sildenafil, which demonstrated that while administering both offered benefits in terms of reducing pulmonary artery remodeling and improving right ventricular function, combination therapy did not show a more pronounced effect in rats, suggesting that there is no synergistic effects between the drugs [22]. This is hypothesized to be related to the overlap between dapagliflozin and sildenafil mechanisms in certain pathways, such as endothelial function, or even due to methodological factors, including dosage and duration. This, however, warrants deeper exploration of SGLT2 inhibitors’ full potential in combinational therapies.

### 4.4 Safety profile

The safety of SGLT2 inhibitors in preclinical PAH models was generally favorable. While only a subset of included studies systematically reported safety outcomes, none reported significant adverse effects related to SGLT2i treatment. Among the included animal studies, survival outcomes were reported in four studies[8,21,22,25]. Notably, the study by Li et al.[8], which demonstrated an opposing effect on pulmonary pressure, reported no significant difference in mortality between groups, whereas the remaining three studies[21,22,25] suggested a survival benefit associated with SGLT2i treatment.

Metabolic parameters such as body weight and glucose levels were also reported. Three studies[23–25] showed a reduction in body weight among animals treated with SGLT2i, aligning with the known weight-lowering effects of this drug class. Two studies[24,25] reported higher glucose readings in treated animals, which may reflect a lower risk of hypoglycemia.

Overall, the safety signals observed in the included studies appear reassuring, with a trend toward improved survival and no significant adverse findings. Nevertheless, the variability in reported outcomes underscores the importance of context-specific safety evaluations in PAH.

### 4.5 Strengths and limitations

This meta-analysis has several strengths. The low heterogeneity across most key outcomes strengthens confidence in the consistency of SGLT2 inhibitors’ effects in PAH animal models. By evaluating multiple outcome domains, including hemodynamics, remodeling, and cardiac function, the study provides a comprehensive assessment of the observed associations. Subgroup analyses by species and drug type demonstrated consistency in the direction of effects across models, supporting the robustness of the findings.

Nonetheless, important limitations must be acknowledged. The overall number of included studies was small (n = 9), with certain outcomes like mean pulmonary artery pressure (mPAP) informed by only three studies, limiting statistical power and the precision of effect estimates. While heterogeneity was low for most outcomes, moderate heterogeneity in two endpoints, RVSP and TAPSE, suggests the presence of unidentified sources of variation. This variability is likely attributable to methodological and measurement-related factors rather than inconsistent biological effects, as RVSP and TAPSE are particularly sensitive to anesthesia protocols, catheter positioning, and operator-dependent echocardiographic techniques in rodent models. Other experimental variables, such as dosing regimen, treatment duration, and timing of assessment, may also contribute to this heterogeneity. To account for such between-study variability, all analyses were performed using random-effects models, which provide more conservative estimates in the presence of heterogeneity.

In addition, the distribution of included studies across individual SGLT2 inhibitors was uneven, with most studies evaluating dapagliflozin, fewer assessing canagliflozin, and only one investigating empagliflozin. This imbalance reflects the current preclinical literature rather than selective inclusion but limits the ability to draw firm conclusions regarding drug-specific effects. Although subgroup analyses by drug type were performed, these analyses were underpowered, and most outcomes did not demonstrate statistically significant differences between individual agents, supporting the interpretation of a class effect rather than superiority of a specific compound.

The inability to formally assess publication bias due to the limited number of studies leaves room for concern, especially given the tendency for negative preclinical results to go unreported. Furthermore, although all included models reflect features of Group 1 PAH, their applicability to the full spectrum of human disease, such as idiopathic, heritable, or connective tissue disease–associated PAH, remains uncertain.

Despite these limitations, the consistent direction of effect across outcomes and models offers a compelling basis for future preclinical and translational investigations.

### 4.6 Future clinical and research perspectives

Our findings highlight the potential of SGLT2 inhibitors as emerging therapeutic candidates for pulmonary arterial hypertension (PAH); however, these results remain hypothesis-generating and must be interpreted with caution. To enhance the translational value, future animal studies should employ diverse disease models and standardized protocols to better reflect human PAH pathophysiology.

One major challenge in translating preclinical success into clinical efficacy lies in interspecies differences, artificial disease induction methods, and the oversimplification of human PAH in rodent models. Addressing this gap requires careful experimental design, improved reproducibility, and mechanistic studies to clarify how SGLT2i affect pulmonary vascular remodeling and right ventricular function.

Ultimately, translation to human trials should be guided by robust, reproducible preclinical evidence. While early safety signals are reassuring, future studies must evaluate long-term outcomes and ensure consistency across models. By systematically addressing these preclinical gaps, the therapeutic potential of SGLT2 inhibitors in PAH can be rigorously assessed and appropriately validated in clinical settings.

## Abbreviations

PAH: Pulmonary arterial hypertension
SGLT2i: Sodium-glucose cotransporter-2 inhibitor
mPAP: Mean pulmonary artery pressure
RVSP: Right ventricular systolic pressure
RV/LV+S: Right ventricular hypertrophy index (right ventricle / left ventricle + septum)
TAPSE: Tricuspid annular plane systolic excursion
PAAT: Pulmonary artery acceleration time
WMD: Weighted mean difference
CI: Confidence interval

## Statements & Declarations

### Author Contributions

Fares Qubbaj contributed to the conception and design of the study, methodology development, data acquisition, formal analysis, drafting of the manuscript, and critical revision of the manuscript.

Ahmad Saeed contributed to data acquisition, methodology development, data curation, drafting of the manuscript, and critical revision.

Osama Younis contributed to data acquisition, formal analysis, drafting of the manuscript, and critical revision.

Nada Al-Awamleh and Zeid Al-Sharif contributed to data acquisition, methodology development, data curation, drafting of the manuscript, and critical revision.

Qais Shaban and Samia Sulaiman contributed to data acquisition, methodology development, drafting of the manuscript, and critical revision.

Ahmad Turk contributed to study conception and design, supervision, project administration, critical revision of the manuscript, and final approval of the version to be published.

All authors approved the final manuscript and agree to be accountable for all aspects of the work.

### Conflicts of Interests

The authors have nothing to declare.

### Funding

No funds, grants, or other support was received.

### Ethics approval

This systematic review and meta-analysis did not involve any direct experimentation on animals. All data were obtained from previously published studies. Therefore, no ethical approval was required for this work.

## Notes

### Competing Interest Statement

The authors have declared no competing interest.

